## Supplemental materials for "Risk Prediction of RNA Off-Targets of CRISPR Base Editors in Tissue-Specific Transcriptomes Using Language Models"

Kazuki Nakamae, Ph.D.

Genome Editing Innovation Center, Hiroshima University,

3-10-23 Kagamiyama, Higashi-Hiroshima, Hiroshima 739-0046, Japan

Hidemasa Bono, Ph.D.

Graduate School of Integrated Sciences for Life, Hiroshima University,

3-10-23 Kagamiyama, Higashi-Hiroshima, Hiroshima 739-0046, Japan

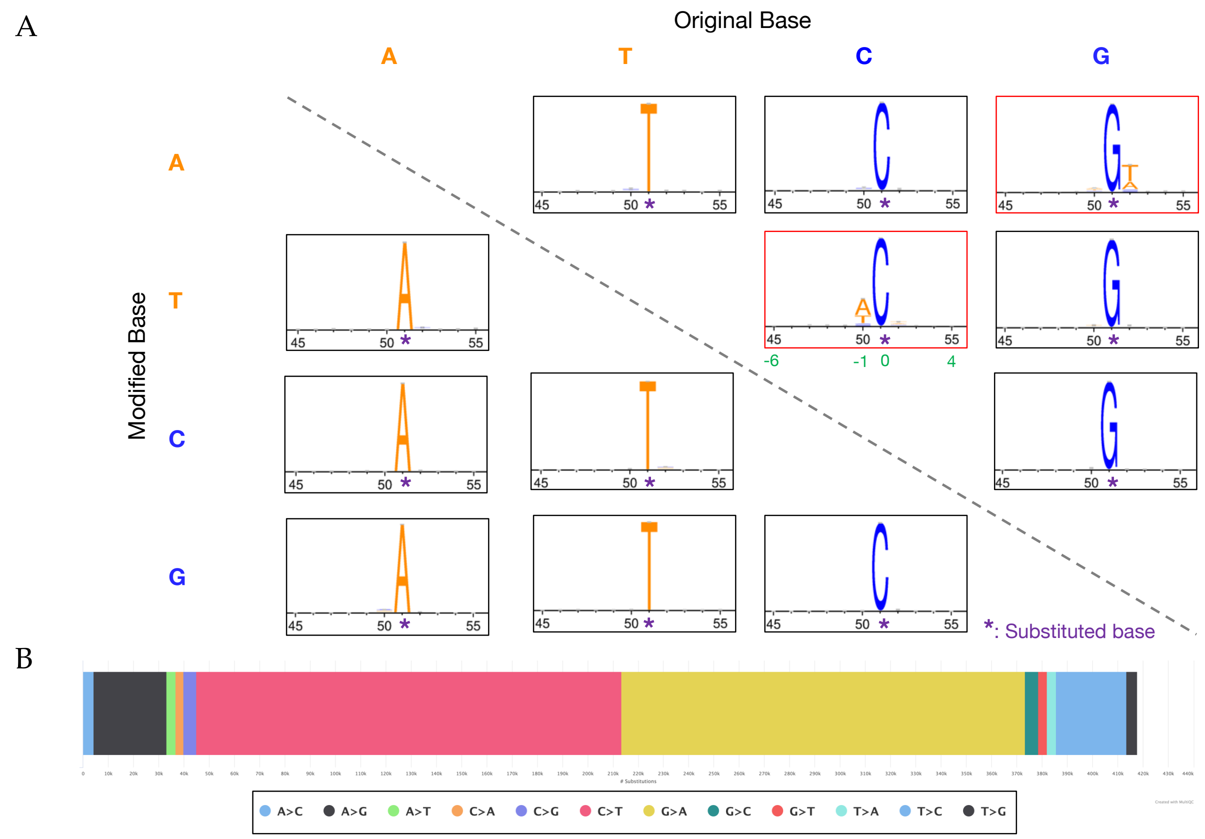


**Supplementary Figure S1.** Overview of substitution patterns and motif preferences identified by PiCTURE. The sequence context and frequency distributions of various substitution classes detected by PiCTURE are shown. (A) The panels are arranged according to the original and substituted nucleotides (columns and rows), with each motif logo representing the flanking sequence preferences (±50 bp) surrounding the substituted bases. The substituted base is located at the relative position of 51 nt (purple asterisk). The red-outlined panels highlight the C-to-T (U) (and complementary G-to-A) substitutions. Black and green numbers indicate distance from 5' -end and substituted base on the horizontal axis, respectively. (B) The bottom bar chart shows the relative proportions of each substitution class, emphasizing the predominance of C-to-T (U) substitutions (G-to-A on the complementary strand) in these datasets.

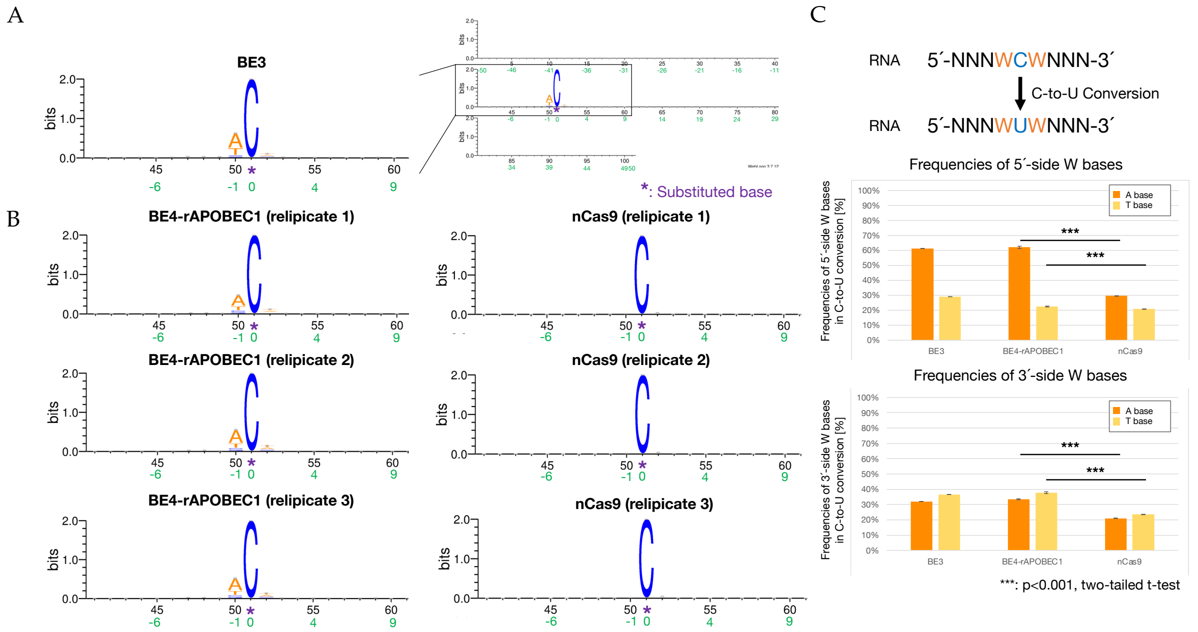


**Supplementary Figure S2.** Motif preferences associated with C-to-U conversions in BE3 and BE4-rAPOBEC1 samples compared to nCas9 negative controls. The results of motif analysis are shown. Sequence logos show that (A) BE3, (B) BE4-rAPOBEC1, and nCas9 samples exhibit distinct motif preferences in C-to-U substitutions. Each motif logo represents the flanking sequence preferences (±50 bp) surrounding the substituted bases. The substituted base is at the relative position of 51 nt (purple asterisk). Black and green numbers indicate distance from 5' -end and substituted base on the horizontal axis, respectively. (C) The bar charts on the right compare the proportions of A and T (U) bases at the 5′ and 3′ neighboring sites of C-to-U conversions among BE3, BE4-rAPOBEC1, and nCas9 samples. Statistical significance is indicated by asterisks (***$p < 0.001$ with two-tailed Welch's t-test).

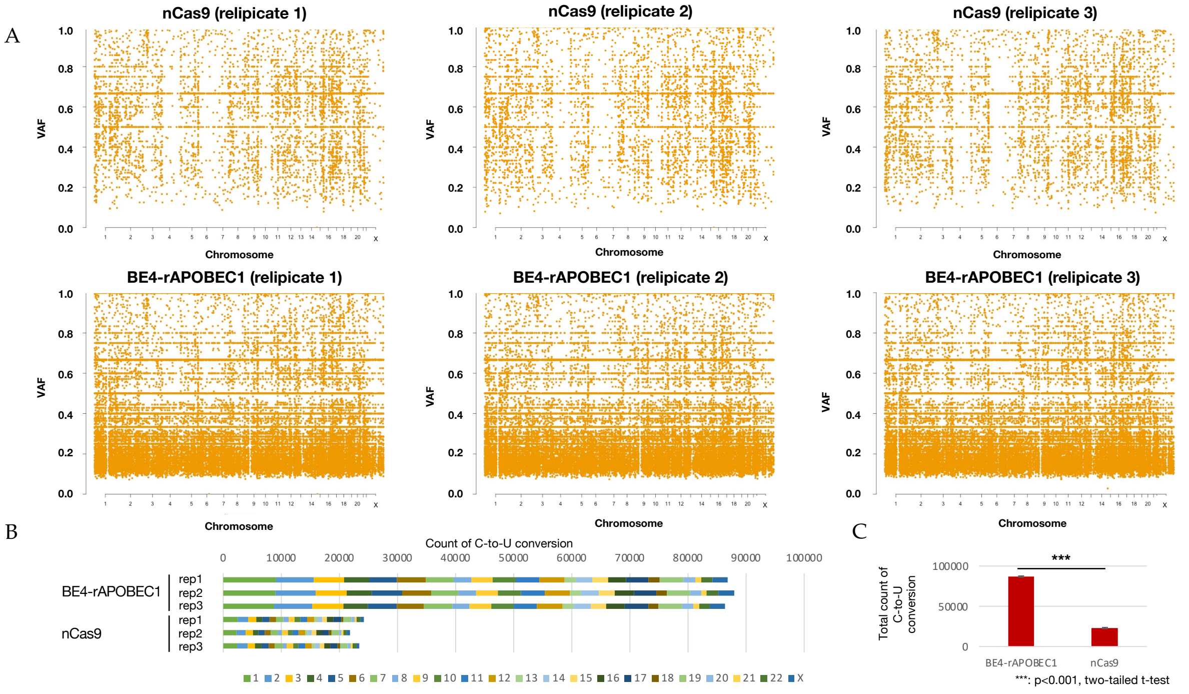

**Supplementary Figure S3.** Chromosomal distribution and abundance of C-to-U conversions in BE4-rAPOBEC1 and nCas9 samples. (A) Dot plots of variant allele frequency (VAF) for C-to-U conversions across individual chromosomes for nCas9 (top row) and BE4-rAPOBEC1 (bottom row) replicates. Each dot represents a genomic region of the transcript with C-to-U conversion, with chromosome positions arranged along the x-axis and VAF along the y-axis. By plotting VAF—the fraction of reads that harbor the C-to-U substitution among all reads mapping to that region—we visualize how frequently the conversion occurs across the dataset. BE4-rAPOBEC1-treated samples show widespread and abundant C-to-U conversions compared to nCas9 controls. (B) Bar chart comparing the relative abundance of C-to-U conversions between BE4-rAPOBEC1 and nCas9 samples per replicate. (C) Bar chart comparing the average total counts of C-to-U conversions with standard error of the mean between BE4-rAPOBEC1 and nCas9 samples. Statistical significance is indicated by asterisks (***$p < 0.001$ with two-tailed Welch's t-test).

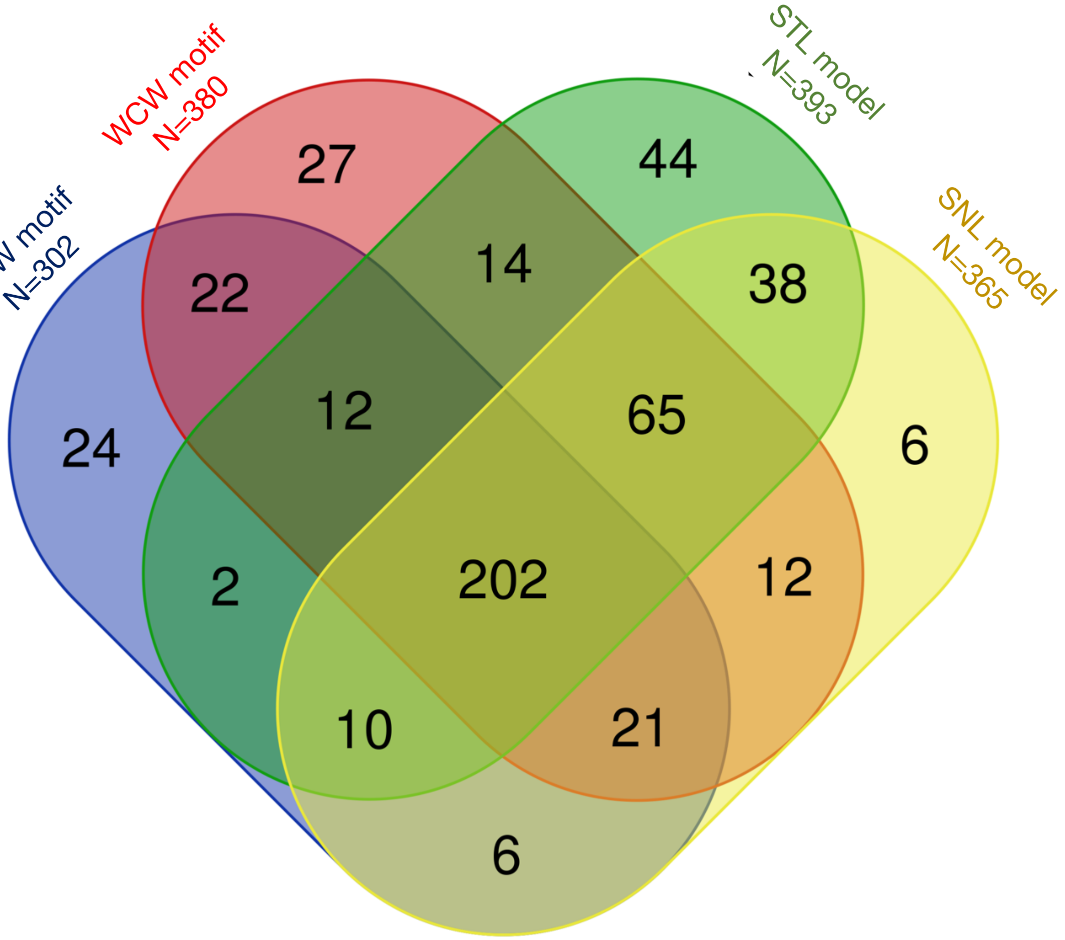


**Supplementary Figure S4.** Overlap of high-risk genes identified by motif-based and machine learning-based classifiers. The Venn diagram compares the high-risk, tissue-specific genes identified by the ACW and WCW motif-based classifiers and the STL and SNL model-based classifiers. Each classifier independently identifies genes susceptible to C-to-U conversions induced by cytosine base editors, with the numbers indicating how many genes are uniquely or commonly detected by each method.

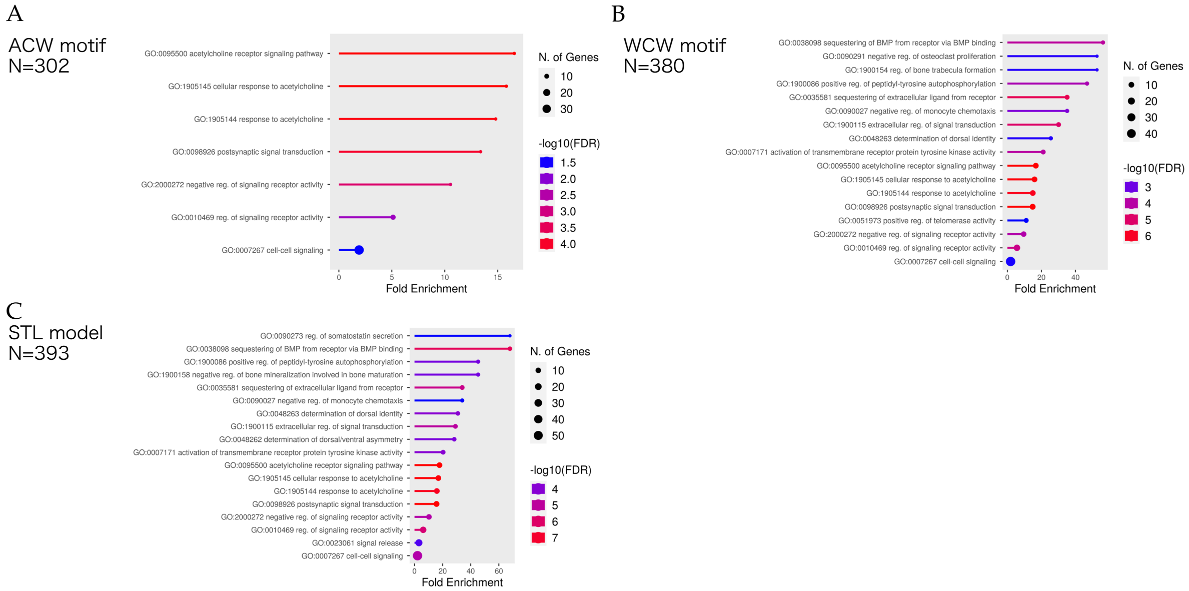


**Supplementary Figure S5.** Gene Ontology (GO) enrichment analyses of high-risk gene subsets identified by different classifiers. The panels present GO Biological Process enrichment analyses for various subsets of high-risk, tissue-specific genes identified by (A) the ACW motif classifier, (B) WCW motif classifier, and (C) STL model. Each bar plot illustrates the enriched GO terms, their fold enrichment values, the number of associated genes, and the statistical significance (–log10(FDR)). N represents the number of high-risk colon-specific transcripts.
